# Deep analysis of the USP18-dependent ISGylome and proteome unveils important roles for USP18 in tumour cell antigenicity and radiosensitivity

**DOI:** 10.1101/2020.03.31.005629

**Authors:** Adan Pinto-Fernandez, Mariolina Salio, Tom Partridge, Jianzhou Chen, George Vere, Helene Greenwood, Cyriel Sebastiaan Olie, Andreas Damianou, Hannah Claire Scott, Henry Jack Pegg, Alessandra Chiarenza, Laura Díaz-Saez, Paul Smith, Claudia Gonzalez-Lopez, Bhavisha Patel, Emma Anderton, Neil Jones, Tim R. Hammonds, Kilian Huber, Ruth Muschel, Persephone Borrow, Vincenzo Cerundolo, Benedikt M. Kessler

## Abstract

The deubiquitylating enzyme USP18 is a major negative regulator of the interferon (IFN) signalling cascade. IFN pathways contribute to resistance to conventional chemotherapy, radiotherapy, and immunotherapy and are often deregulated in cancer. USP18 is the predominant human protease that cleaves interferon-stimulated gene ISG15, a ubiquitin-like protein tightly regulated in the context of innate immunity, from its modified substrate proteins *in vivo*. In this study, using advanced proteomic techniques, we have expanded the USP18-dependent ISGylome and proteome in a chronic myeloid leukaemia (CML)-derived cell line (HAP1) treated with type I IFN. Novel ISGylation targets were characterised that modulate the sensing of innate ligands, antigen presentation and secretion of cytokines. Consequently, CML USP18-deficient cells are more antigenic, driving increased activation of cytotoxic T lymphocytes (CTLs) and are more susceptible to irradiation. Our results suggest USP18 as a pharmacological target in cancer immunotherapy and radiotherapy.

## Introduction

Despite its clinical success in inducing regression of certain tumours, immune-checkpoint blockade (ICB) therapy is limited by being effective in only a fraction of patients. Many patients develop resistance^1^ to treatment; Moreover, ICB frequently has profound side effects, calling for the discovery of predictive biomarkers of ICB success and for novel ways to boost ICB therapy further. Numerous studies point towards defects in interferon-dependent pattern recognition pathways ^2–8^ and the antigen-presentation pathway ^9,10^ as the main resistance mechanisms tumour cells use to avoid the immune system and to escape the effects of immunotherapy ^11, 12^. In addition to the roles of elements of the innate immune response in modulating the efficacy of cancer immunotherapy, it has also been reported that radiotherapy promotes the expression of interferon-stimulated genes (ISGs) that are involved in resistance to ionizing irradiation^5,13–16^. Immunomodulatory imide drugs (IMiDs) such as thalidomide analogues are another category of immunomodulators that are routinely used to treat patients with multiple myeloma and also lead to increased expression of ISGs ^17^. Therefore, there is strong evidence pointing at the innate immune response as a resistance mechanism that cancer cells use to survive the effects of immune checkpoint blockade and other cancer therapies.

One of the main regulators of the innate immune response is the deISGylating enzyme USP18 (Fig.1a). Despite its full name (ubiquitin-specific protease 18), USP18 is not a deubiquitylating enzyme (DUB). In fact, it is the predominant protease described to date to recognise and to remove specifically the ubiquitin-like protein ISG15 from modified proteins *in vivo*. USP2, USP5, USP13, USP14, and USP21 DUBs are also active in ISG15 *in vitro* assays, but they do not seem to deconjugate ISG15 in a cellular context ^18–22^. ISG15 is structurally very similar to a dimer of ubiquitin, and its expression is, like that of USP18, tightly regulated by type I IFN. ISG15 is not conserved across species and its functions may vary between organisms. It has been identified only in vertebrates; moreover, human and murine ISG15 protein sequences only share 64% and 74% homology and similarity, respectively ^23,24^. ISG15-null patients present with inflammation due to deregulation of the type I interferon response and, in contrast to observations made in to mice, ISG15 is not involved in susceptibility to viral infection ^25,26^. USP18 deficiency sensitises murine cells and mice to interferon or to activators of this pathway^27^. USP18 functions that are independent of ISG15 seem to be crucial in mice, since the phenotype observed in *Usp18-/-* animals, including brain inflammation and de-regulated STAT1 signalling, is not rescued by removing *Isg15* ^28^. In humans, however, free ISG15 stabilises USP18 and also acts as a negative regulator of the IFN pathway ^26,29^.

**Figure 1:**
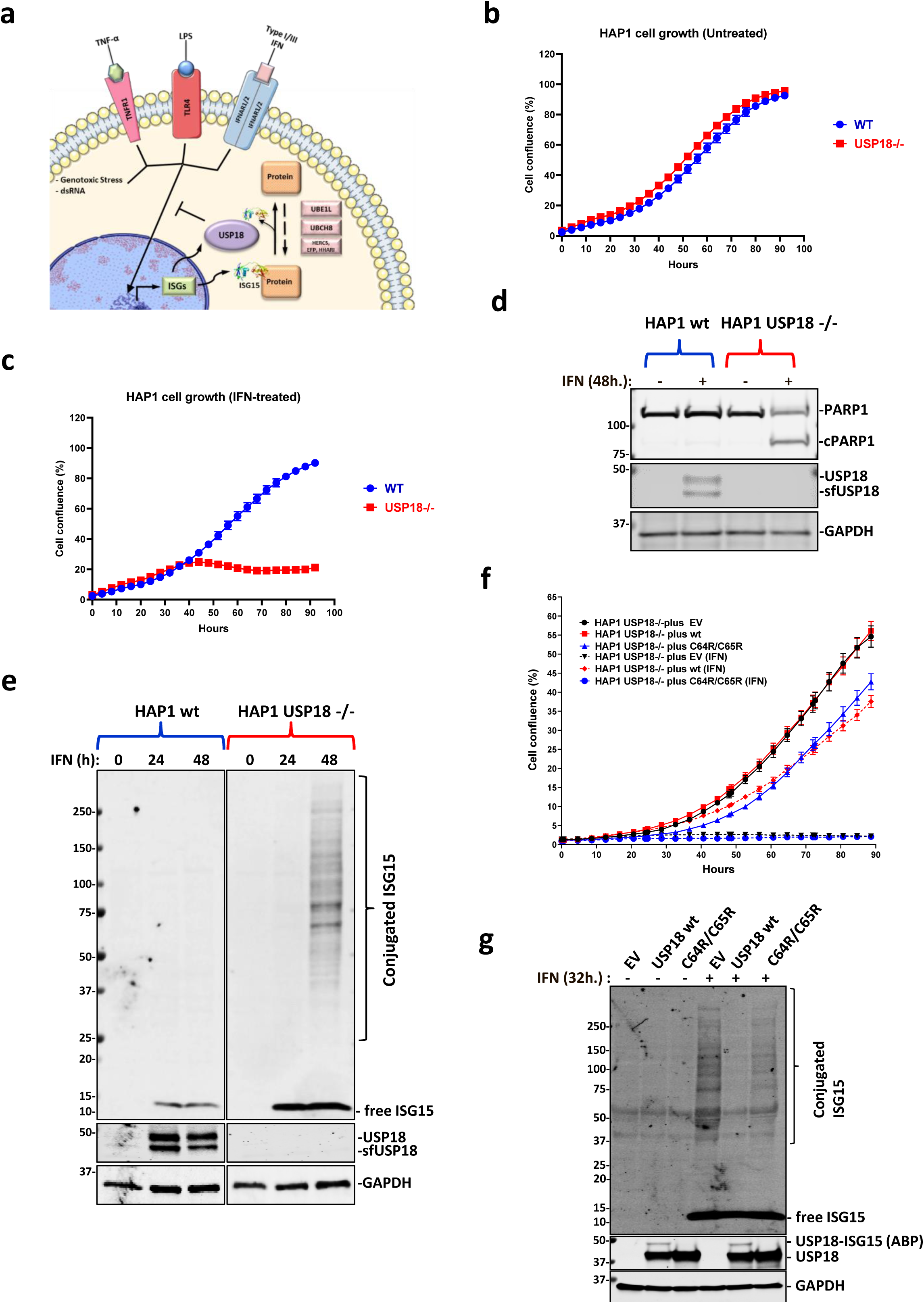
USP18 alters protein ISGylation and cell viability in presence of IFN. **a)** Depiction of the roles of USP18. USP18 is an interferon-stimulated gene (ISG) that acts as a negative regulator of the type I interferon (IFN) signalling pathway and removes ISG15, another ISG structurally similar to a dimer of ubiquitin, from modified proteins. **b)** HAP1 CML-derived wild-type (WT) and USP18 knockout cells (KO) growth is comparable. **c)** The growth of HAP1 cells deficient for USP18 is inhibited by type I IFN treatment. **d)** Immunoblot showing cleavage of the pro-apoptotic marker cPARP in HAP1 USP18-/-cells after 48 h. treatment with IFN. **e)** Immunoblot showing induction of USP18 after 24 h. and 48 h. of treatment with IFN in WT cells and accumulation of conjugated ISG15 in USP18-deficient cells in the same conditions. **f)** Stable re-expression of USP18 WT, but not of a catalytically inactive mutant USP18 C64R/C65R, in USP18-deficient cells rescue them from the growth-inhibitory effects of IFN. **g)** Immunoblot of cell extracts from USP18 KO cells expressing USP18 WT and USP18 C64R/C65R. Ectopic expression of WT but not catalytically inactive USP18 prevents the accumulation of ISGylated proteins upon IF treatment (top panel). ISG15 propargylamide (ISG15-PA) activity-based probe (ABP) labels only USP18 WT, whilst it is unreactive towards the catalytically inactive mutant (middle panel). GAPDH was used as a loading control in all the immunoblots.

Both ISG15 and USP18 contribute to innate immune responses and are also important in other cellular processes such as autophagy and protein translation ^30,31^. USP18 has been linked to cancer development, being overexpressed in lung, colon, pancreas and breast cancer, but also associated with autoimmune diseases, infection and neuroinflammation ^22,32–34^. *Usp18-/-* mice present a strong accumulation of ISG15 conjugates in tissues as well as a prolonged increase in STAT1-dependent activation ^35^. These mice are hypersensitive to IFN and Poly I:C (a synthetic double-stranded RNA) stimulation, and are more resistant to intracerebral infection. In some mice genetic backgrounds, *Usp18* deficiency has also been linked to brain abnormalities ^22,27,35^. Supporting its role as a negative regulator of immunogenicity, it has been reported that silencing USP18 also increases surface expression of peptide-loaded MHC-II ^36^. Notably, USP18 has also been described to have a role as a resistance factor to traditional chemotherapy agents Bortezomib and Mafosfamide in tumour cells ^37,38^, suggesting that its inhibition may serve as a boost for such therapies.

Protein ISGylation profiles in human cells have been previously reported, but with limited depth and predominantly based on overexpression systems ^39–44^. To obtain deep coverage and modified residue information in a more physiologically relevant context, we decided to combine advanced proteomic techniques with the immunoaffinity purification of GlyGly tryptic peptides ^45,46^ in wild-type (WT) and USP18-/- knockout (KO) HAP1 cells, in order to analyse the first USP18-dependent ISGylome of human cancer cells. We uncovered novel ISGylated substrates that are controlled by USP18, further emphasizing its role as a master regulator of the innate immune response. Our results provide strong evidence for USP18 in regulating antigenicity and radiosensitivity, highlighting its potential as a cancer target.

## Results

### USP18 catalytic activity is essential for cell survival in the presence of type I IFN

We decided to study the effects of type I interferon Alpha 2 (IFNa2; IFN hereafter) on human HAP1 cells, a chronic myeloid leukaemia (CML)-derived cell line, in the presence or absence of the USP18 gene. Under normal culture conditions, both cell lines proliferate at the same rate (Fig. 1b). However, while the parental cells are completely insensitive to treatment with 1000 U/mL of IFN (Fig. 1c), the USP18-deficient cells stop growing after 24h (Fig. 1c and Fig. S1a) and start to die as visualised by microscope phase-contrast imaging Fig. S1b. Supporting this phenotype, a molecular indicator of programmed cell death, the cleavage of the protein PARP, is only observed in the KO cells treated with IFN (Fig. 1d). Consistent with previous literature ^22,27^, deletion of USP18 also led to the accumulation of ISG15 conjugates in our cellular model, shown as a strong smear after 48 hours of treatment with IFN (Fig. 1e) and an increase in ISG15 levels and STAT1 phosphorylation in immunofluorescence analysis after 24 hours of IFN stimulation (Fig. S1c). Strikingly, the accumulation of ISGylated proteins is barely noticeable in HAP1 WT cells after IFN treatment (Fig. 1e).

USP18 functions as a negative regulator of the IFN pathway have been linked not only to its ability to remove ISG15 from proteins, but also to inhibitory protein-protein interactions with IFNAR2 and STAT2 ^47^. In order to evaluate the importance of USP18 catalytic activity in our model, we re-expressed wild-type (WT) and a catalytically inactive mutant, where we mutated the catalytic cysteine and a second adjacent cysteine (C64R/C65R) in order to get a fully inactive enzyme, in HAP1 USP18 KO cells. Stable transfectants expressing the WT protein rescued KO cells from the toxic effects of IFN (Fig. 1f and Fig. S1d) and also prevented the accumulation of ISGylated proteins (Fig. 1g). Unexpectedly, and in contrast with what has been published in murine systems, re-expression of USP18 C64R/C65R did not rescue the KO cells from IFN toxicity (Fig. 1f and Fig. S1d) or from the accumulation of ISGylated proteins (Fig. 1g). As a control to evaluate the activity of the two proteins, we performed deISGylating activity assays using ISG15 activity-based probes that bind irreversibly to catalytically active USP18 and allow the visualization of the active enzyme as an increase in the molecular weight ~15 kDa (USP18 plus ISG15 probe) by immunoblotting ^18,48^. We observed that only wild-type USP18 is able to bind the activity-based probe, whereas the C64R/C65R mutant is completely inactive (Fig. 1g, middle panel). As a control for inhibition of the IFN-receptor by direct binding of WT or C64R/C65R USP18, we performed immunoblots against the ISGs HERC5, IFIT3 and ISG15. Both, WT and C64R/C65R USP18 are able to inhibit the expression of these three ISGs, although the inhibitory effect seems to be smaller for the catalytically inactive mutant (Fig. S1e), perhaps pointing at a slightly impaired ability to bind IFNAR2 or STAT2 of this mutant due to structural changes induced by the mutations.

### USP18-dependent GlyGly peptidome/proteome reveals tumour cell ISGylome

Digestion with trypsin of proteins modified either with ubiquitin, ISG15 or NEDD8 leaves a unique GlyGly motif on the modified lysine (Fig. 2a and 2b). Enriching for GlyGly-modified peptides followed by mass spectrometry (MS) analysis provides access to information on not only the prevalence/amount of the modification but also on its exact position in the protein sequence. A recent report exploited this to analyse cellular ISGylomes in mice, although using a different genetic cellular background strategy to the one used here and in the context of infection ^49^. In this study, we employed a similar approach to conduct the first comprehensive analysis of the USP18-dependent ISGylome in human cancer cells.

**Figure 2:**
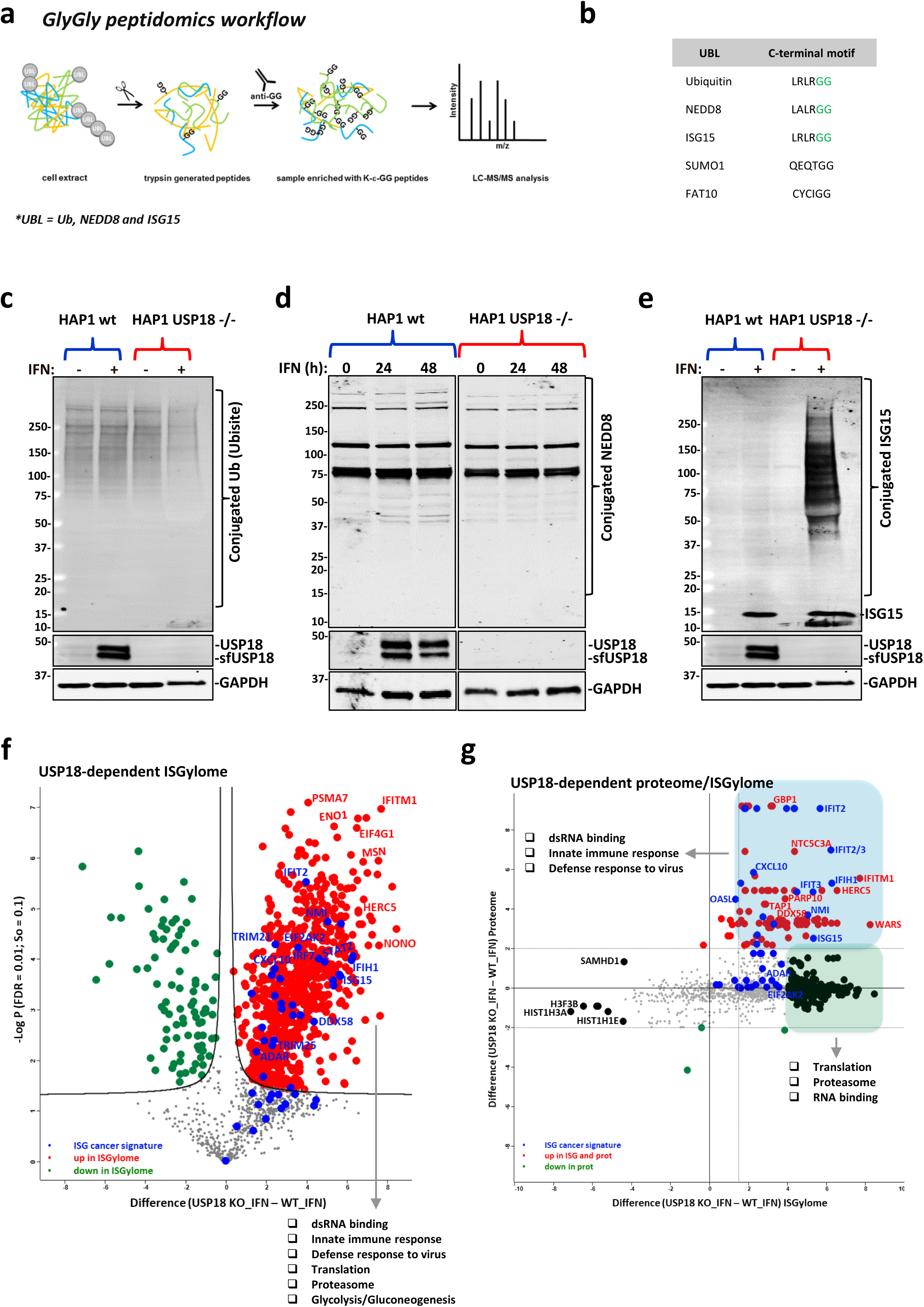
USP18 is the major cellular de-ISGylase. **a)** Workflow for analysis of GlyGly-modified peptides. **b)** Comparative table of the C-terminal motifs of the different ubiquitin-like proteins. ISG15, Ubiquitin, and NEDD8 leave a unique GlyGly motif (in green) after trypsin digestion of modified proteins. **c)** Ubiquitin-conjugated and **d)** NEDD8-conjugated proteins do not increase after stimulation with type I interferon (IFN) in USP18 KO cells. **e)** ISG15 modification is accumulated in the same conditions as in c). **f)** Comparative volcano plot of the GlyGly modified peptides in HAP1 WT and USP18 KO cells treated with IFN for 48 h showing a strong and significant USP18-dependent upregulation of GlyGly-modified peptides in the KO cells treated with IFN for 48 h. In red are shown the upregulated peptides in the KO cells, in green the upregulated peptides in WT cells, and in blue the modified peptides from an ISG-cancer signature ^8^ (the statistical cut-off values used for all the proteomic analyses performed in this study are FDR: 0.01 and s0: 0.1). Pathway enrichment analysis showed the upregulated peptides in the USP18 KO cells come from proteins involved in the indicated biological processes. **g)** Scatter plot of the cross-comparative analysis of ISGylome and proteome in the same conditions as f). In red are shown the upregulated ISGylated proteins in the KO cells, in green the upregulated modified proteins in WT cells, and in blue the modified proteins from an ISG-cancer signature ^8^.

Even though GlyGly peptidomics is not specific for ISG15, we hypothesised that KO of USP18 (the main deISGylating enzyme *in vivo*) in cells should lead to accumulation of ISGylated substrates upon type I IFN stimulation that would be suitable for identification using GlyGly immunoaffinity purification of the generated tryptic peptides. To validate our hypothesis, we first wanted to verify, by immunoblotting, that ubiquitylation and NEDDylation were not affected across our four experimental conditions (HAP1 WT and USP18 KO in the presence/absence of IFN). As shown in Fig. 2c and 2d, ubiquitylation, and NEDDylation did not seem to be affected, if something, we saw a small reduction of ubiquitylated proteins in the USP18 KO cells treated with IFN. In contrast, IFN treatment of the KO cells led to a massive accumulation of conjugated ISG15 material (Fig. 2e). Based on these observations, we would expect to detect mainly the accumulated ISGylated proteins when applying the GlyGly peptidomics in these particular experimental conditions. 2,341 GlyGly-modified peptides were identified in this mass spectrometry-based experiment, corresponding to 2,172 protein groups. As the volcano plot in Fig. S2a shows, there are no significant changes in the comparative GlyGly peptidomes of HAP1 cells treated with and without IFN, consistent with the results from the immunoblots. However, and again in line with the gel results, following treatment of USP18 KO cells with IFN and comparison of their GlyGly peptidome to that of the WT cells (also treated with IFN), we observed a massive USP18-dependent, increase in GlyGly peptides, corresponding to 476 proteins, in the KO cells (Fig. 2f). After gene ontology/pathway enrichment analysis, we could reinforce the role of USP18 as a major regulator of the innate immune response with significant enrichment of ISG15-modified peptides for dsRNA binding and closely related proteins such as ADAR, DDX58 (RIG-I), DDX60, DHX58, OAS1/2, EIF2KA2 (PKR), IFIH1, DDX3X, DHX9, STAT1 and others that are part of innate immunity pathways. Also, we observed a strong upregulation of ISGylated (immuno)-proteasomal subunits PSMA7, PSMB9, PSMB10, and PSME2 as well as TAP1, all of which are part of the antigen presentation pathway. Proteins involved in translation and glucose metabolism processes were also represented with numerous ISGylated protein members (Fig. 2f). Interestingly, we also found that proteins from a recently published signature of interferon-stimulated genes (ISGs) expressed constitutively in a subset of cancer cell lines ^8^ were also strongly upregulated and represented in our ISGylome (Fig. 2f). The global proteome of the same samples was analysed in parallel (Fig. S2b, 2c, and 2d). USP18 is expressed in very low amounts when the IFN pathway is inactive and therefore, unsurprisingly, we did not detect major changes in either the proteome (Fig. S2c) or the GlyGly-peptidome (Fig. S2g) of WT cells compared to USP18 KO cells growing in unstimulated conditions. However, when the IFN pathway was activated in WT cells, we could see an enrichment of ISGs and elements from the above-mentioned cancer ISG signature (Fig. S2d). In agreement with the ISGylome data, the USP18-dependent proteome (or ‘interferome’) in the presence of IFN showed an even stronger upregulation of ISGs, mainly consisting of dsRNA-related enzymes, proteasome subunits, and IFITs, indicating a general activation of the interferon pathway (Fig. S2b, e, and f). Interestingly, many of the upregulated proteins in the USP18-dependent ‘interferome’ overlap with a subset of factors recently identified to be upregulated in metastatic melanoma responders to ICB ^50^. In particular, USP18 regulates the expression and/or ISGylation status of PSME1, PSMB9, PSMB10, B2M, TAPBP, TAP1, TAP2, CBR3, STAT1, IFIT1, GBP1, GBP2, ERAP1 and MAGE, all linked to better responses to ICB therapy with PD1 inhibitors (Table S1). In line with previous literature ^51–53^, when cross-comparing the USP18-dependent ISGylome and proteome (in IFN-treated cells) data sets, we could observe that in our case ISGylation is not leading to a decrease in protein levels (Fig. 2g), in contrast to ubiquitylation. However, a strong regulatory ISGylation of ribosomal proteins, proteasomal subunits, and some RNA-related enzymes was observed without affecting their expression, (Fig. 2g; in green) and also an accumulation of modified proteins, mainly ISGs (Fig. 2g; in blue). The latter effect may putatively not be due to ISGylation leading to a stabilisation of proteins, but might instead reflect the roles of USP18 as an inhibitor of late-stage IFN signalling. However, the potential role of ISG15 as stabiliser of a certain set of ISGs cannot be excluded at this point ^53^.

### USP18 controls ISGylation of the dsRNA modifier ADAR (p150) and the innate ligand sensor PKR and regulates their activity

To verify our ISG discovery data, we used immunoblotting with specific antibodies against a group of selected ISGylated hits such us ADAR, PKR, HERC5, and IFIT3, and we could observe a modified version of these proteins (consistent with, at least, an increase in the MW of ~15 kDa) in the KO cells treated with IFN, but not in the other conditions (Fig. 3a and 3d). To further validate ISGylation as the observed modification, we silenced the ISG15 gene using specific siRNA sequences in the USP18-deficient cells and we treated them with IFN. As shown before, the dsRNA regulating enzyme ADAR (p150) was modified in the KO cells in the presence of IFN. However, after partial silencing of ISG15, the modification became significantly reduced (Fig. 3b). As the ultimate validation technique for our GlyGly-based ISGylome, we decided to perform a classic interactome analysis after immunoaffinity purification (IAP-MS) of endogenous ISG15 proteins in the same experimental conditions. After optimisation of the immunoprecipitation (IP) conditions, we could efficiently purify ISG15 and its conjugated substrates (Fig. 3c). Again, we did not observe an increase of ubiquitylated proteins in the inputs, and we did not co-immunoprecipitate ubiquitin or ubiquitylated proteins together with ISG15 (Fig. 3c, lower panel). In the same IP eluates, we observed an enrichment of modified HERC5, ADAR, PKR and IFIT3, all ISGylated in the GlyGly data, in the USP18 KO cells treated with IFN (Fig. 3d). Mass spectrometry analysis of the eluates showed enrichment of very similar proteins to the matching GlyGly peptidome in the USP18 KO cells treated with IFN (Fig. 3e). From a total of 312 ISGylated proteins found to interact with ISG15 in a USP18-dependent manner, 110 proteins overlapped when comparing the two techniques (Fig. S3a), primarily the ones showing a stronger enrichment in the volcano plot in Fig. 3e (highlighted and labelled in green) and in the comparative scatter (ISG15 interactome *versus* GlyGly peptidome) plot in Fig. 3f, upper right area. These complementary experiments validated our approach for analysis of the site-specific ISGylome using GlyGly peptidomics.

**Figure 3:**
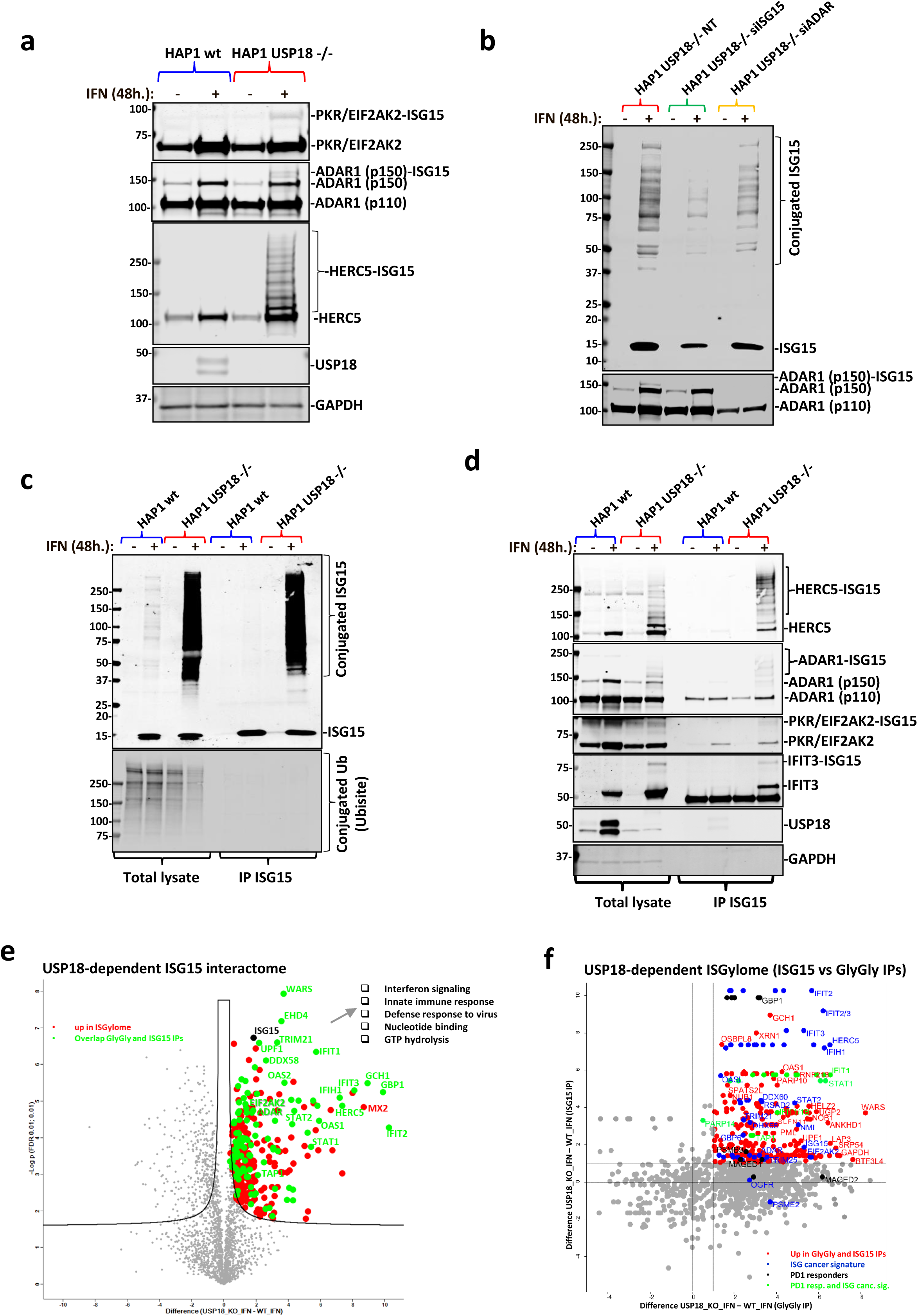
USP18-deletion exacerbates the cellular ISGylated protein network. **a)** Immunoblot using specific antibodies against the indicated proteins showing additional ISGylation of these proteins in the HAP1 USP18 KO cells treated with IFN. **b)** HAP1 USP18 KO cells were transfected with siRNA sequences targeting ISG15 and ADAR and treated with IFN for the indicated times. After ISG15 silencing, modified ADAR (ADAR1 (p150)-ISG15; lower panel) is no longer visible by immunoblot. **c)** HAP1 WT and USP18 KO cells were treated with IFN for the indicated times, lysed and subjected to immunoprecipitation (IP) with ISG15 antibodies. Enriched ISGylated proteins were present in eluates from the USP18 KO cells (top panel) but there was no enrichment for ubiquitylated proteins (lower panel). **d)** Immunoblot of the same eluates in c) against the indicated proteins showing enrichment for ISGylated species. **e)** Volcano plot of comparative proteomic analysis of the IP eluates in c) and d). Pathways enriched are shown. Comparative scatter plot **f)** of the GlyGly peptide IP and the ISG15 IP, performed in the same experimental conditions showing a strong overlap between the two data sets and strong enrichment of members of the ISG cancer gene signature ^8^ and factors up-regulated in PD1-responders ^50^.

The double-stranded RNA-specific adenosine deaminase ADAR (adenosine deaminase acting on RNA) is an essential gene for tumour cells that are positive for the above-mentioned signature of ISGs ^8^ and for a subset of tumour cell lines with increased expression of ISGs ^54^. Even more importantly, ADAR has been shown to be involved in resistance to ICB therapy and irradiation in preclinical models ^7^. The molecular basis of these observations is to the excessive accumulation of dsRNA induced by the absence of ADAR and the subsequent activation of the sensors for this innate ligand, mainly PKR and the axis DDX58/IFIH1, leading to cell death and tumour inflammation ^7,8,54^. As we found ADAR to be highly modified in the KO cells treated with IFN, we decided to check if in our cellular model ADAR is also essential upon activation of the IFN pathway. To do so, we silenced ADAR with specific siRNA sequences in HAP1 WT cells and we subsequently treated them with IFN. In line with what has been published, and in a similar fashion to USP18 depletion, ADAR genetic inactivation sensitises HAP1 cells to IFN (Fig. 4a). In addition to its effects on cell growth, we also evaluated the activation of the dsRNA sensor PKR, measured by immunoblotting using an antibody against PKR when phosphorylated at threonine 446, in WT and USP18 KO cells treated with IFN for 0, 6, 24 and 48 hours. Both, ADAR and PKR are ISGylated upon IFN treatment but only in the USP18 KO cells, which is particularly noticeable in PKR’s case since it was modified after just 6h of treatment (Fig. 4b). ISGylation of PKR fits with its markedly higher phosphorylation in the KO cells compared with a relatively modest activation in the WT cells. ISGylation of PKR (although on different residues) has been previously reported to activate PKR ^52^, which aligns well with our observations. We identified 7 lysine residues in the sequence of ADAR that are potentially ISGylated (K433, K637, K763, K781, K798, K895, and K996; Fig. 4c). When highlighting these residues in the structure of the deaminase domain of ADAR2 bound to RNA, three of them seem to directly contribute to the binding of the enzyme to RNA (K637, K781, and K996; Fig. 4d). Finally, we measured the levels of the innate ligand dsRNA in our cells using specific antibodies against dsRNA by immunofluorescence as an indication of ADAR activity. ADAR enzymes are responsible for binding to double-stranded RNA (dsRNA) and converting adenosine (A) to inosine (I) by deamination, thus reducing the intracellular levels of dsRNA. The results shown in Fig. 4e and 4f illustrate the accumulation of dsRNA in a time-dependent manner only in the USP18 KO cells treated with IFN for 0, 24 and 48 hours. This accumulation matches well with the ISGylation levels of ADAR p150 seen in Fig. 4b and points at USP18 and ISG15 as key regulators of ADAR deaminase activity.

**Figure 4:**
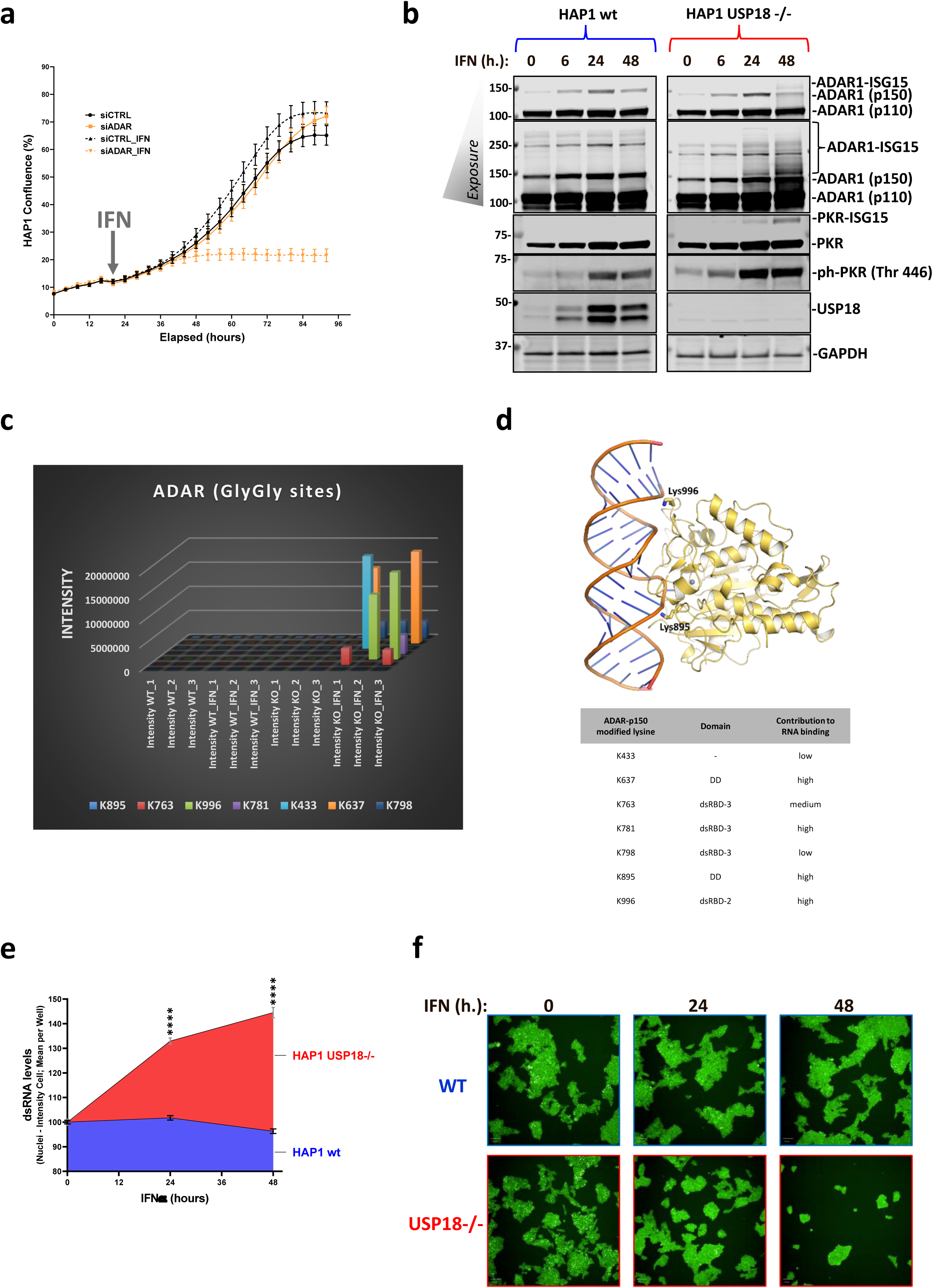
USP18-dependent ISGylation of ADAR leads to enzymatic activity inhibition and dsRNA accumulation. ADAR knockdown using siRNA renders HAP1 WT sensitive to growth inhibition by type I IFN as shown by the growth curves in **a)**. **b)** Immunoblots of a time-course experiment showing an increase in modified ADAR and PKR and activation of PKR (phosphorylated PKR in Thr 446) USP18-deficient cells treated with IFN for the indicated times. **c)** A plot of the ADAR GlyGly-modified lysines and their ion intensities measured by MS. **d)** dsRNA bound ADAR2 deaminase domain structure (PDB code 5ED2). Lysine residues corresponding to the ADAR1 GlyGly sites are represented with sticks. The Zn^2+^ cation is represented as a grey sphere (top panel). Table showing the ADAR1 potential GlyGly sites indicating their domain location and their contribution to RNA binding. A score of 1 indicates the highest contribution and 3 the lowest and away from the RNA binding pocket. DD = deaminase domain. dsRBD = dsRNA binding domain (lower panel). **e) and f)** Analysis of the dsRNA levels after IFN treatment in WT and KO cells by immunofluorescence using specific antibodies (n=6 per condition).

### USP18 regulates MHC class I antigen presentation, PD-L1 expression and stimulation of a T cell functional response

In the USP18-dependent ISGylome and proteome of cancer cells treated with type I IFN, we observed ISGylated components of the antigen presentation pathway (Fig. 2e, 3e). To test whether USP18 modulates antigen presentation and T cell stimulation, we pulsed the HLA-A2 positive HAP1 cell line (WT and USP18 KO; with and without pre-treatment with IFN for 24 hours) with different concentrations of the HLA-A2-restricted Melan-A_26-35_ peptide and then co-cultured them with Melan-A specific CTL for 18 hours ^55–57^.

Peptide-pulsed WT or USP18 KO HAP1 cells (+/- type I IFN pre-treatment) stimulated phenotypic activation of T cells (as assessed by upregulation of the activation markers CD25 and CD137) with equal efficiency (Fig.5a). However, both with and without IFN pre-treatment, the USP18 KO cells triggered a more robust functional response by the T cells, as evidenced by greater T cell production of IFNγ following peptide recognition (Fig. 5b). Since the strength of the activating stimulus received by T cells determines the nature of their response, with phenotypic activation and production of chemokines such as MIP1β being more readily triggered than production of cytokines such as IFNγ ^58^, this suggests that USP18-deficient tumour cells exhibit superior antigenicity. Non-peptide-pulsed HAP1 USP18 KO cells expressed a similar level of HLA-A2 to the parental cells and HLA-A2 was equivalently upregulated on both following exposure to type 1 IFN (Fig. 5c, top left histogram plot). HLA-A2 was further upregulated on both WT and KO cells after pulsing with the Melan-A26-35 peptide and co-culture with T cells, likely in response to IFNγ secreted by the T cells (Fig. 5c; top right and lower histogram plots). However a proportion of the peptide-pulsed cells exhibited downregulation of HLA-A2 to levels below those expressed on non-peptide-pulsed cells following co-culture with T cells, potentially as a consequence of HLA-A2 internalisation and/or degradation (Fig. 5c; top right and lower histogram plots). Notably, high levels of HLA-A2 expression were retained on a greater proportion of the USP18 KO cells than their WT counterparts (Fig. 5c; bottom graph), putatively contributing to their greater antigenicity.

**Figure 5.**
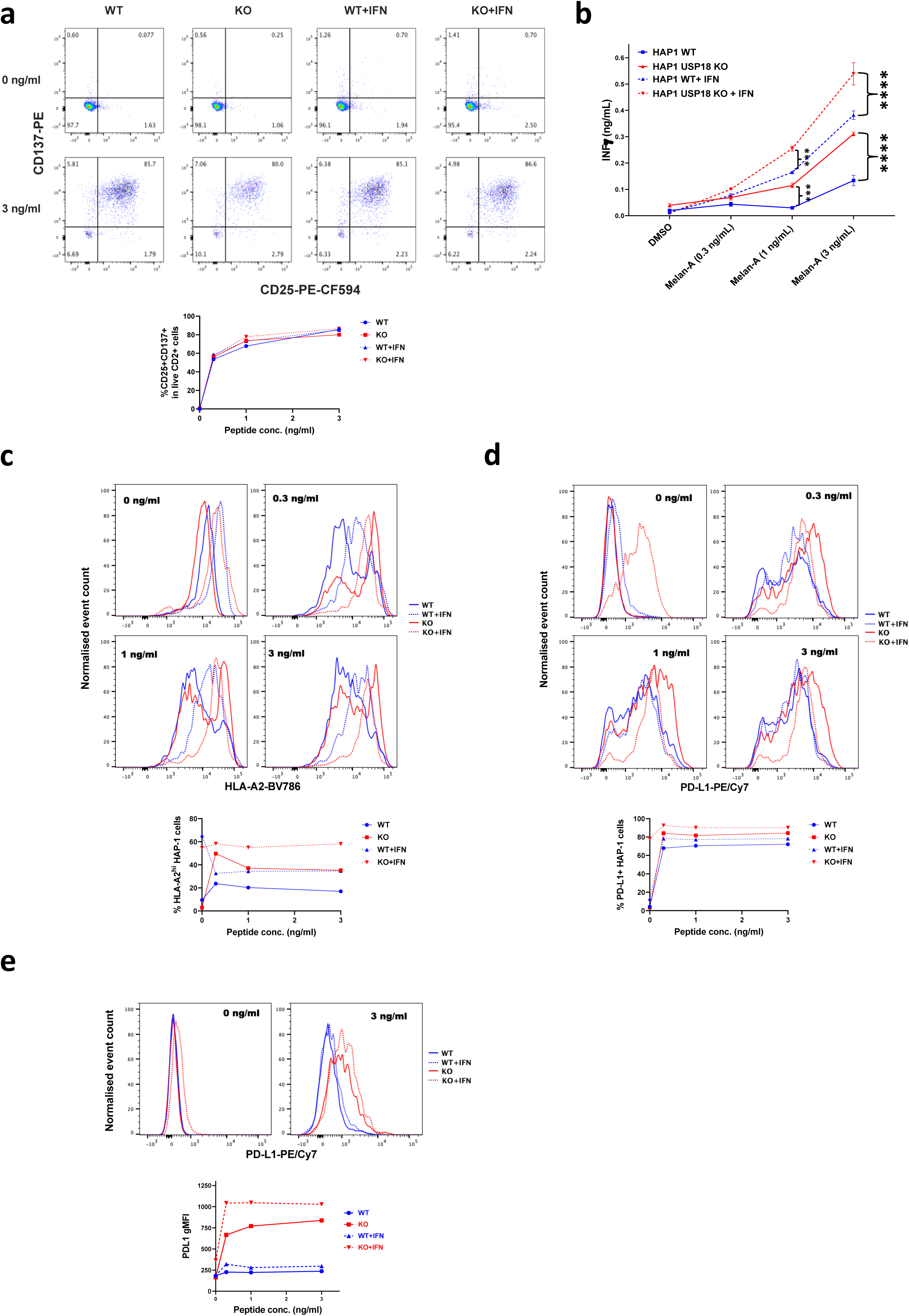
USP18 regulates major histocompatibility complex class I (MHC-I) antigen presentation, PD-L1 expression and stimulation of a T cell response. Wild-type or USP18-deficient HAP1 cells were pre-treated with or without IFN-a, then pulsed with Melan-A26-35 peptide at the indicated concentrations and co-cultured with Melan-A-specific T cells. **a)** Upregulation of activation markers on T cells was assessed by flow cytometry. Dotplots show representative examples of CD25 and CD137 expression on live CD2^+^ cells (T cells) following co-culture with the indicated peptide-pulsed HAP1 cells and the proportion of T cells expressing both CD25 and CD137 is plotted below. **b)** T cell release of IFNy into HAP1-T cell co-culture supernatants was measured by ELISA. Bars show mean values from 3 replicates and for the statistical analysis, we applied two-way ANOVA tests including multiple comparison testing via the Dunnett method available through the GraphPad Prism software. P value style is GraphPad: NS, P = 0.1234, *P = 0.0332, **P = 0.0021, ***P = 0.0002, ****P < 0.0001. Expression of **c)** HLA-A2 and D) PD-L1 on HAP1 cells following co-culture with T cells was measured by flow cytometry. Histogram plots of the data are shown above, and the plots below show the percentage of cells c) expressing high levels of HLA-A2 (defined as greater than those expressed on non-peptide-pulsed cells not pre-treated with type 1 IFN and **d)** expressing PD-L1. **e)** PD-L1 expression on T cells following co-culture with the indicated HAP1 cells. Representative histogram plots are shown above, and the geometric mean fluorescence intensity (gMFI) of PD-L1 expression on T cells from each co-culture condition is plotted below. The results shown in this figure are representative of data from 1 of 3 experiments and the FACS shows information of three replicates pooled into one.

Strikingly we observed a strong upregulation of PD-L1 on both tumour cells (Fig. 5d) and T cells when USP18 was absent (Fig. 5e). This effect was antigen dependent, although the KO cells pre-treated with type I IFN presented a high basal expression of PD-L1 in the absence of peptide. PD-L1 protein expression in tumours has been described as a potential predictive biomarker for sensitivity to immune checkpoint blockade (ICB) with PD1/PD-L1 inhibitors^59^. Also, we observed an upregulation of PD-L1 levels on the T cells when co-cultured with USP18 KO cancer cells, regardless of treatment. In contrast, very little upregulation of PD-L1 was noted on T cells stimulated with WT cells (+/- IFN), after addition of peptide (Fig. 5e). PD-L1 is expressed on activated T cells and is required for T cell conditioning and dendritic cell maturation ^60^. However, its expression on T cells has been mostly linked to inhibition of their responses ^61,62^. Other studies indicates that expression of PD-L1 on CD8^+^ and CD4^+^ T cells correlates with patient response to ICB, suggesting PD-L1 expression in CD8^+^ T cells as a prognostic marker in melanoma ^12,63,64^.

### USP18 deficiency sensitises cancer cells to radiation therapy (RT)

Efficacy of local radiotherapy (RT) has been linked to the ability of the irradiated cells to repair lethal DNA damage but also to the induction of T cell-dependent cell killing. This immune attack has been linked to the induction of the type I IFN response after tumour RT ^13,15,16,65^ and also seems to be ADAR-dependent^7^. As shown above, USP18-decient HAP1 cells displayed greater cellular death following activation of IFN signalling by exposure to type I IFN in tissue culture. Thus, we hypothesized that loss of USP18 in cancer cell could also augment their sensitivity to ionizing radiation. Indeed, USP18 KO cells exhibited a significantly lower clonogenic survival rate after radiation in tissue culture compared with WT (representative images in Fig. 6a and summary in Fig. 6b). In addition, the average colony diameter of USP18 KO cells was smaller than that of WT (Fig. 6c) indicating that USP18 deficiency also results in greater growth delay after exposure to ionizing radiation. Consistently, following a single dose of 5Gy irradiation, the cell number ratio of WT over USP18 KO cell increased over time (Fig. 6d).

**Figure 6:**
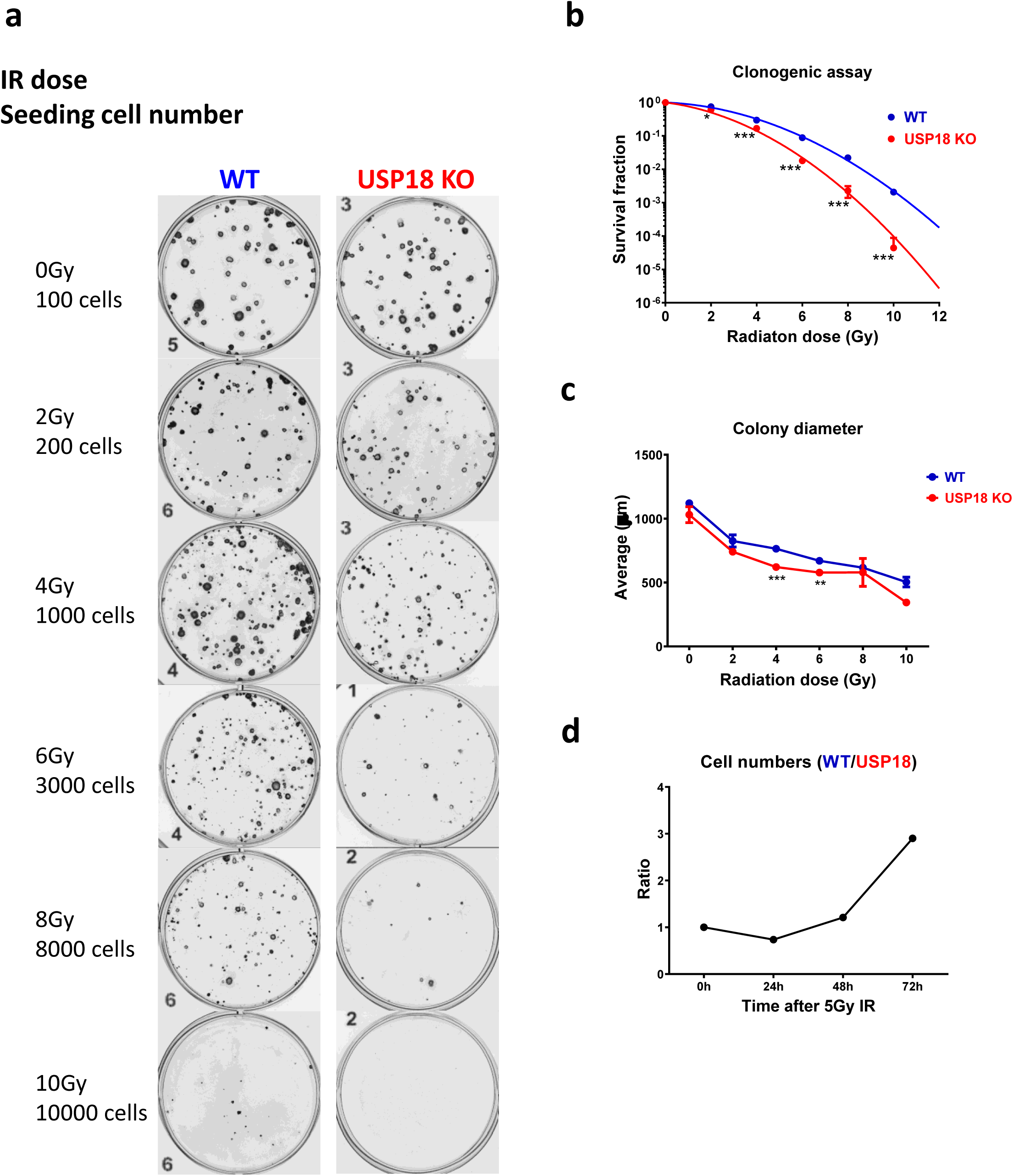
Deletion of USP18 increases cellular radiosensitivity (IR): **a)** Representative images of clonogenic assay with HAP1 WT cells (left column) and HAP1 USP18KO (right column). The IR doses and seeding cell numbers were indicated. **b)** Quantitation of data from the clonogenic assay measured as survival fraction in the same conditions as in a). n = 3 biological repeats. **c)** The diameter of the counted colonies in a) and b). n =3. **d)** Cell number ratio of WT vs USP18 -/- HAP1 cells at indicated time points after a single dose of 5Gy IR. Data represent mean ± SD. Comparison of two means was performed by the one-way ANOVA (* P < 0.05, ** P< 0.01, *** P< 0.001).

## Discussion

This study provides a comprehensive profile of the USP18-regulated human endogenous ISGylome at a site-specific resolution. USP18 is the predominant deISGylating enzyme that processes ISG15 *in vivo* and, like ISG15, its expression is induced in the presence of agonists of the interferon pathway. USP18-deficient cells are sensitive to IFN treatment and strongly accumulate ISGylated proteins without noticeable effects on ubiquitylation or NEDDylation. This effect seems to be dependent on the catalytic activity of USP18, contradicting some previously reported studies on mice and humans. Notably, in this study, we used specific ISG15 activity-based probes to demonstrate that the catalytically inactive mutant enzyme (C64A/C65A) is actually inactive. This is particularly important in the case of USP18 since there is an additional cysteine (C65) adjacent to the catalytic cysteine (C64). When performing activity assays, we found that single mutants C64S and C64R were still active (data not shown). Here, we were able to identify 2,341 GlyGly sites and 476 enriched modified proteins that, after exhaustive validation using orthogonal techniques, we were confident to label them as ISGylated proteins and map their modified sites. When comparing our data with previous ISGylome studies, 537 new ISGylated human proteins were identified (Fig. S3b). Hence, this study expands considerably the human, USP18-dependent, ISGylome, suggesting novel regulatory cellular functions by ISGylation. We used different bioinformatics tools such as Weblogo in order to find consensus sequences for ISG but without success. We cannot exclude that applying artificial intelligence-based analysis could help to identify the sequences more susceptible to this modification ^66^. A cross-comparison between the GlyGly peptidome data and the matching proteome further confirmed that ISGylation is not involved in protein degradation, but rather stabilisation, perhaps through co-translational modification ^67^. Also, the overall expression level of many of the ISGylated proteins does not change, reinforcing the role of ISG15 as a PTM modulating enzymatic activities ^52,68^.

Pathway analysis could identify some significantly enriched biological processes modulated by USP18-dependent ISGylation such as dsRNA binding, innate immune response, defence to virus, translation and glycolysis/gluconeogenesis. Many of these proteins are ISGs, and a number of them are part of a signature of genes that are upregulated in a subset of tumour cells, independent of immune infiltration ^8^ ISG-positive cancer cells are sensitive to ADAR loss, another ISG with deaminase activity, controlling the levels of dsRNA, an innate ligand ^8,54^ We identified USP18-dependent ISG15ylation in seven lysine residues for ADAR (K433, K637, K763, K781, K798, K895, and K996), localised within the dsRNA binding (dsRBD) and deaminase domains (DD), which may potentially affect both enzymatic activity and substrate binding affinity. ADAR has been recently described as a major factor for resistance to immunotherapy and radiotherapy ^7^. In our model, ADAR gene silencing using siRNA sequences sensitises cancer cells to IFN stimulation in a similar fashion as USP18 does. We could detect ISGylated ADAR by immunoblotting in USP18-deficient cells after 24 hours treatment with IFN, and this time point matches the time when the growth of the cells starts to be inhibited by this cytokine. The described effects of ADAR loss in overcoming immunotherapy and radiotherapy are linked to its ability to reduce the levels of innate ligand dsRNA and subsequent inhibition of the dsRNA sensors PKR, and MDA5/RIG-I, involved in growth inhibition and tumour inflammation, respectively ^7,8,54^ ISGylation of PKR has been reported to activate its activity, and we could also see ISGylation and activation of PKR in the USP18 KO cells treated with IFN by immunoblotting. Our data, therefore, suggested that USP18-dependent ISGylation of ADAR was inhibiting its activities, inducing the accumulation of dsRNA, resulting in further activation of PKR. Immunofluorescence analysis using specific dsRNA antibodies showed a significant and time-dependent accumulation of dsRNA in the USP18 KO cells after IFN treatment, indicating that ADAR activity is inhibited by USP18-dependent ISGylation under these conditions.

In addition to ADAR, PKR, RIG-I, and MDA5, we found other proteins involved in antigen presentation and resistance to immunotherapy, such as TAP1, GBP1, STAT1, IFIT1, PSMB10, PSMB9, GBP2, MAGE and PARP14 ^50^, also regulated by USP18-dependent ISGylation. Many of these genes are part of processing MHC class I antigens, suggesting that the USP18 KO cells could possibly be more antigenic. Indeed, our results confirmed that USP18 KO cells had an enhanced ability to stimulate IFNγ secretion by Melan-A antigen-specific T cells, which was associated with elevated HLA-A2 expression levels. Interestingly, PD-L1 levels, a prognostic factor for ICB therapy when using PD1 inhibitors, were also higher in USP18 KO as compared to WT cells, and PD-L1 was also upregulated on the T cells following interaction with these cells. This is noteworthy as other studies have linked the expression of PD-L1 on CD8^+^ and CD4^+^ T cells with patient response to ICB, suggesting PD-L1 expression in CD8^+^ T cells as a prognostic marker in melanoma ^12,63,64^. As the innate immune response is activated by radiotherapy ^13,15,16^ and ADAR recently described as an important factor in resistance to RT ^7^, we decided to perform a second round of functional experiments consisting of irradiating our cancer cells and evaluating viability and clonogenicity after RT. The results clearly suggest that USP18 deficiency sensitises tumour cells to RT in a dose-dependent manner, most likely by interfering with end-stage IFN signalling.

Directing the immune system against tumours, also known as cancer immunotherapy, is supplementing or replacing more conventional forms of therapy as the first line of treatment for many tumour types due to striking curative effects in a subset of the treated patients. However, resistance mechanisms prevent this therapy from being 100% effective. The innate immune response has been described as one of the main resistance mechanisms against resistance to ICB and the discovery of factors involved in reactivation of the innate response, such as ADAR, has been a priority for many academic labs and also for pharma. So far, inhibition of ADAR has been unfruitful. Our data suggest that USP18 inhibition represents a novel strategy to block ADAR functions, to activate the immunoproteasome/antigen presentation machinery and innate ligand sensors. As a consequence, this may help to overcome resistance to immunotherapy and RT, potentially turning ‘cold’ tumours into ‘hot’ tumours.”

## Acknowledgments

We would like to thank the TDI MS Laboratory/Discovery Proteomics Facility for their technical support and Sebastian Nijman for providing the USP18-deficient HAP1 cells. Work in the BMK lab was funded by FORMA Therapeutics, by EPSRC grant EP/N034295/1 and by the Chinese Academy of Medical Sciences (CAMS) Innovation Fund for Medical Science (CIFMS), China (grant number: 2018-I2M-2-002).

## Methods

### Cell lines and T cells

Chronic myeloid leukaemia (CML)-derived HAP1 wild-type (WT) cells and HAP1 USP18-/- (KO) were a kind gift from the laboratory of Sebastian Nijman and were cultured in IMDM media (Gibco #12440-53) supplemented with 10% FBS (Gibco #10500-64) at 37 °C in a humidified 5% CO2 atmosphere.

Melan-A specific CTL lines were isolated from healthy blood donors as described ^57^. Briefly, dendritic cells pulsed with Melan A peptide (ELA_26-35_ ELAGIGILTV, Sigma) were incubated with autologous PBMC; after 12 days melan-A specific CD8 T cells were sorted with HLA-A2-ELA class I tetramers and expanded as described ^57^.

### Stimulation of cells

Cells were treated with 1000 U/mL of human interferon alpha 2 (Alpha2b) from PBL Assay Science (Cat. No. 11105-1) for the indicated times.

### DNA plasmids and generation of stable cell lines

The Flag-HA-USP18WT construct was purchased from Addgene (#22572) and the Flag-HA-USP18C64R/C65R construct was generated using the Q5^®^ Site-Directed Mutagenesis Kit (NEB, E0554S) and the primers OL87: TGGACAGACCcggcggCTTAACTCCTTG and OL88: ATGTTGTGTAAACCAACC. Primer were designed using the NEBaseChanger online tool: http://nebasechanger.neb.com/ and the successful clone confirmed by sequencing (Eurofins).

Transfected HAP1 cells were selected, pooled, and subsequently cultured in growth media supplemented with puromycin (Gibco) at 1 μg/mL.

### siRNA reagents

ISG15 siRNA on-TARGET plus SMART plus (Dharmacon #L-004235-03-0005) and ADAR siRNA on-TARGET plus SMART plus (Dharmacon #L-008630-00-0005).

### Cell transfection

USP18 cDNA plasmids were transfected with Lipofectamine LTX and Plus (Invitrogen #15338-100) following the manufacturer’s instructions. The indicated siRNA sequences were transfected with Lipofectamine RNAimax (Invitrogen #13778-150) following the manufacturer’s instructions.

### Antibodies

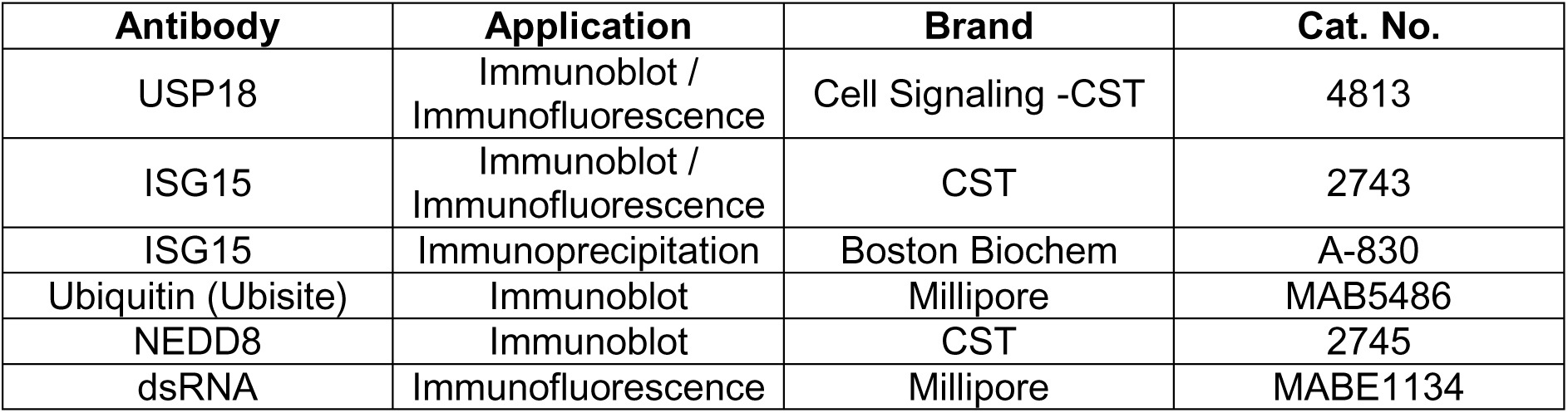
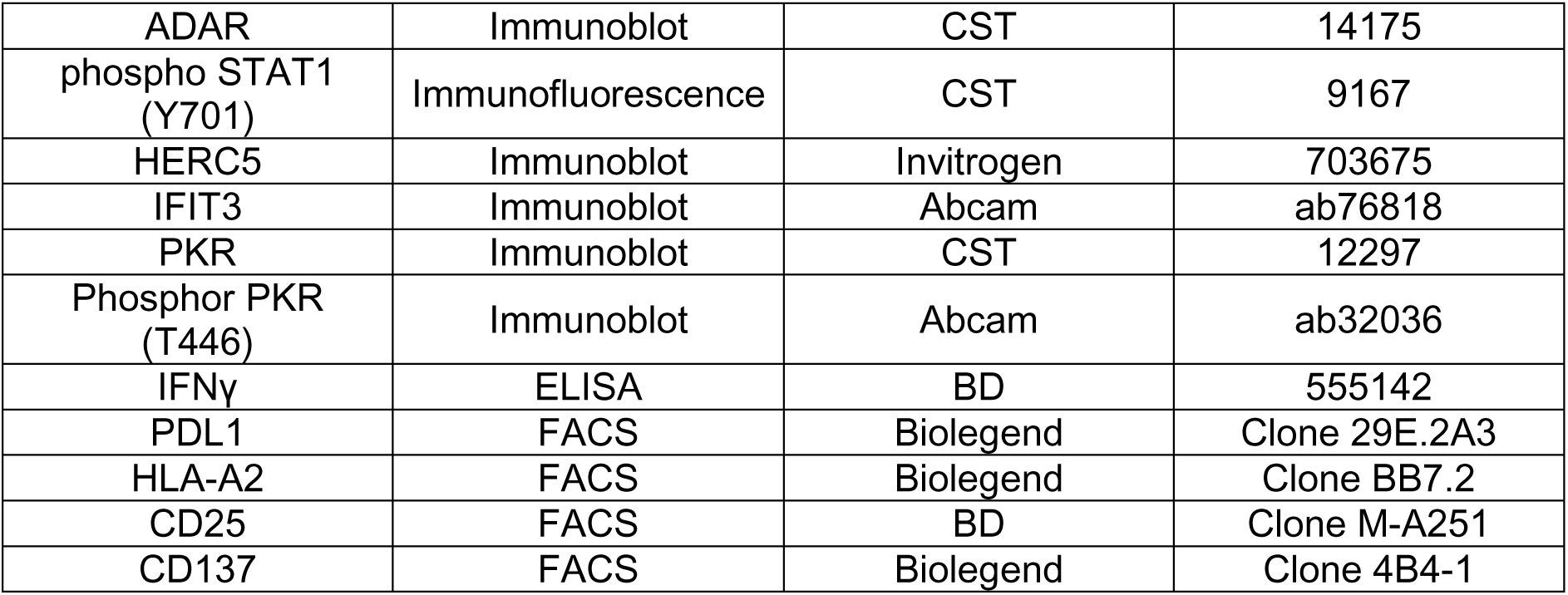

### Western blotting

Cells were washed with ice-cold PBS and lysed with NP-40 lysis buffer (50 mM Tris pH 7.4, 0.5% (v/v) NP40, 150 mM NaCl and 20 mM MgCl2) buffer containing protease and phosphatase inhibitors. For Western blotting, 25 μg of protein was then fractionated on Tris–glycine SDS-PAGE gradient (4–15% acrylamide) gels (BioRad; #3450123), transferred onto PVDF membranes (Millipore; IPFL00010), and detected with the indicated antibodies using a LI-COR detection system.

### USP18 ABP

ISG15-PA was synthesised in the Kessler lab using the ISG15 C-term (79-156) construct from Addgene (#110760) and DUB ABP assays were performed as previously described^69^. Briefly, HAP1 cells were lysed in glass beads lysis buffer (GBL: 50 mM Tris, pH 7.5, 5 mM MgCl2, 0.5 mM EDTA, and 250 mM Sucrose) and the extracts were labelled with ISG15-PA for 45 min at 37°C, separated in 4-15% SDS-PAGE Tris-Glycine gradient gels (BioRad; #3450123) and immunoblotted with specific anti-USP18 antibodies.

### Cell growth, morphology and proliferation assays

10000 HAP1 cells were seeded in 6-well plates and cell were imaged in an IncuCyte Zoom Imager (Essen Bioscience) for live-cell imaging during the indicated times at the indicated intervals the phase contrast channel. Cell growth was determined by measuring the percent confluence over time.

### Immunofluorescence microscopy

HAP1 control or HAP1 USP18 KO cells were seeded in a 96 well plate and left overnight to overcome stress. Cells were then treated with 1000 U/mL of human interferon alpha 2 (Alpha2b) for the indicated times. The cells were washed with PBS and fixed using 4% paraformaldehyde (sc-281692, ChemCruz) for 10 minutes. The cells were permeabilised using 100% methanol for 10 minutes at −20 °C. The cells were then blocked with 5 % BSA in PBS for one hour at room temperature. The primary antibodies were diluted in PBST (PBS + 0.05 % Tween20) in the following concentrations: USP18 (1:100), ISG15 (1:100) and pSTAT1 (1:400). The primary antibodies were incubated overnight at 4 °C. Cells were then washed with PBST and subsequently incubated with secondary antibody mix consisting of Goat anti-rabbit conjugated alexa-647 (1:10000) and 4’,6’-diamidino-2-phenylindole (Dapi) (2.5 μg/mL), for 1 hour at room temperature. The cells were then imaged using the Opera Phenix (Perkin Elmer).

For dsRNA analysis, the same workflow was used with the exception of the cells, that were fixed and permeabilised using Cytofix/Cytoperm buffers (BD biosciences), following the manufacturer’s instructions. dsRNA antibody was used at a 1:100 dilution and the secondary donkey anti-mouse at a 1:500 dilution.

### Identification of ISGylated proteins using GlyGly peptidomics and matching proteomics

HAP1 lysates were used for GlyGly immunoprecipitation using PTMScan Ubiquitin Remnant Motif Kit (Cell Signaling), according to manufacturer’s protocol ^45,46^. Briefly, 20 mg of extracts were solubilized and denatured in 10 mL lysis buffer (20 mM HEPES, pH 8.0, 9 M urea, 1 mM sodium orthovanadate, 2.5 mM sodium pyrophosphate, 1 mM β-glycerophosphate), reduced using dithiothreitol (4.5 mM final) for 30 minutes at 55 °C. This was followed by alkylation using iodoacetamide (100 mM final) for 15 minutes at room temperature in the dark. Samples were subsequently diluted fourfold in 20 mM HEPES, pH 8.0 (~2 M urea final), followed by digestion with trypsin-TPCK (Worthington, LS003744, 10 mg/mL final) overnight at room temperature. Samples were then acidified using trifluoroacetic acid (1 % final), and desalted using C-18 Sep-Pak (Waters) cartridges according to the manufacturer’s protocol. At this point, 20 μg of digested protein were separated for matching proteome control analysis. Peptides were lyophilized and re-suspended in 1.4 mL immunoprecipitation IAP buffer (PTMScan), and the remaining insoluble material cleared by centrifugation and anti-GlyGly antibody beads added followed by rotation and kept 4°C for 2 hours. Beads were subsequently washed twice using 1 mL IAP buffer, followed by three water washes. Immunoprecipitated material was eluted twice in 55 and 50 μL 0.15 % trifluoroacetic acid in water. Peptide material was desalted and concentrated using C-18 Sep-Pak (Waters) cartridges according to the manufacturer’s protocol. Purified GlyGly-modified peptide eluates and matching proteome material were dried by vacuum centrifugation, and re-suspended in buffer A (98 % MilliQ-H20, 2 % CH_3_CN and 0.1 % TFA).

### ISG15 interactome immunoprecipitation

HAP1 cells were lysed with Co-IP lysis buffer (20 mM Hepes pH 8.0, 150 mM NaCl, 0.2 % NP-40, 10 % Glycerol, 5 mM NEM, phosphor and protease inhibitor cocktails (25 x 10^6^cells per condition) and subjected to immunoprecipitation using 5 μg ISG15 antibodies (Boston Biochem #A-380) plus 25 μL of protein G Sepharose slurry (Invitrogen; #15920-10) for 16 hours at 4 °C. Beads were washed 4 times with Co-IP lysis buffer and immunoclomplexes were eluted with 2X Laemmli. 10 % of the eluates was used for immunoblotting with the indicated antibodies. The remaining eluate was prepared for MS analysis as previously described ^69,70^. Briefly: immunoprecipitated sample eluates were diluted to 175 μ L with ultra-pure water and reduced with 5 μL of DTT (200 mM in 0.1 M Tris, pH 7.8) for 30 min at 37 °C. Samples were alkylated with 20 μL of iodoacetamide (100 mM in 0.1 M Tris, pH 7.8) for 15 min at room temperature (protected from light), followed by protein precipitation using a double methanol/chloroform extraction method. Protein samples were treated with 600 μ L of methanol, 150 μL of chloroform, and 450 μL of water, followed by vigorous vortexing. Samples were centrifuged at 17,000 g for 3 min, and the resultant upper aqueous phase was removed. Proteins were pelleted following the addition of 450 μ L of methanol and centrifugation at 17,000 g for 6 min. The supernatant was removed, and the extraction process was repeated. Following the second extraction process, precipitated proteins were re-suspended in 50 μL of 6 M urea and diluted to <1 M urea with 250 μ L of 20 mM HEPES (pH 8.0) buffer. Protein digestion was carried out by adding trypsin (from a 1 mg/ml stock in 1 mM HCl) to a ratio 1:100, rocking at 12 rpm and room temperature overnight. Following digestion, samples were acidified to 1 % trifluoroacetic acid and desalted on C18 solid-phase extraction cartridges (SEP-PAK plus, Waters), dried, and re-suspended in 2 % acetonitrile and 0.1 % formic acid for analysis by LC-MS/MS as described below.

### Liquid chromatography-tandem mass spectrometry (LC-MS/MS) analysis

LC-MS/MS analysis was performed using a Dionex Ultimate 3000 nano-ultra high-pressure reverse-phase chromatography coupled on-line to a Q Exactive HF (GlyGly), Fusion Lumos (ISG15 interactome) or a Q Exactive (Matching proteome) mass spectrometer (Thermo Scientific) as described previously ^69–71^. In brief, samples were separated on an EASY-Spray PepMap RSLC C18 column (500 mm × 75 μm, 2 μm particle size, Thermo Scientific) over a 60 min (120 min in the case of the matching proteome) gradient of 2–35 % acetonitrile in 5 % dimethyl sulfoxide (DMSO), 0.1 % formic acid at 250 nL/min. MS1 scans were acquired at a resolution of 60,000 at 200 m/z and the top 12 most abundant precursor ions were selected for high collision dissociation (HCD) fragmentation.

### Data analysis

From raw MS files, searches against the UniProtKB human sequence data base (92,954 entries) and label-free quantitation were performed using MaxQuant Software (v1.5.5.1). Search parameters include carbamidomethyl (C) as a fixed modification, oxidation (M) and deamidation (NQ as variable modifications, maximum 2 missed cleavages (3 for the GlyGly peptidome analysis), matching between runs, and LFQ quantitation was performed using unique peptides. Label-free interaction data analysis was performed using Perseus (v1.6.0.2), and volcano and scatter plots were generated using a t-test with permutation FDR = 0.01 for multiple-test correction and s0 = 0.1 as cut-off parameters.

Other graphs were generated using GraphPad PRISM 8 and Excel and for the statistical analysis, we applied two-way ANOVA tests including multiple comparison testing via the Dunnett method available through the GraphPad Prism software. P value style is GraphPad: NS, P = 0.1234, *P = 0.0332, **P = 0.0021, ***P = 0.0002, ****P <0.0001.

Gene ontology and pathway enrichment analysis were performed using STRINGdb (https://string-db.org/cgi/input.pl?sessionId=9clBKQVOpdjY&input_page_show_search=on) and g:PROFILER (https://biit.cs.ut.ee/gprofiler/gost).

Flow cytometry data were analysed with Flowjo 10, upon gating on live singlets.

### FACS/T cell activation pulsing with Melan-A peptides

50000 HAP1 WT and USP18-/- cells (+/- IFNα-2a, 1000 U/mL; 24h.) were plated in 96-well plates (flat bottom, Costar) and pulsed with the indicated amounts of Melan-A peptides (ELAGIGILTV, Sigma) for 2 hours at 37C. Cells were washed three times with RPMI media (Sigma) prior to addition of melan A-specific T cells (healthy donor-derived) at a ratio 2 tumour cells: 1 immune cell for 18 hours. T cell activation (FACS) and cytokine release in the supernatant (ELISA) were analysed subsequently.

For FACS analysis, triplicate wells were poo led, cells were stained 20min at room temperature with a near infrared live/dead marker (Biolegend) and then with a cocktail of titrated antibodies, 30min on ice. Samples were acquired on a BD Symphony flow cytometer. Flow cytometry data were analysed with Flowjo 10, upon gating on live singlets.

### ELISA

Enhanced ELISA plates (Corning) were coated with diluted anti-IFNγ antibodies (2 μg/mL) in coating buffer (100 mM NaHCO3) and left overnight at 4 °C. Plates were washed 6 times with PBS / 0.01 % Tween and blocked with PBS / 10 % FCS for 2 hours at room temperature. Diluted standards and samples in PBS / 10 % FCS were added to the plate and incubated for 3 hours at 37C. Plates were washed 6 times with PBS / 0.01 % Tween prior addition of biotinylated anti-cytokine detecting mAb (2 μg/mL in PBS / 10 % FCS) for 45 minutes at room temperature. Plates were then washed 8 times and avidin-peroxidase (2.5 μg/mL in PBS / 10 % FCS. Sigma) was added for 30 minutes. Finally, plates were washed 10 times with PBS/Tween and substrate was added (o-phenylenediamine dihydrochloride tablets) and then quenched with 0.2 N sulfuric acid (Sigma) after colour appears visible. Plates were read at OD 490 nm in a microplate reader.

### Clonogenic assays after RT

Cancer cells were seeded in 6-well plates and treated with a range of IR doses (D): 0-10 Gy. The colonies were stained and assessed on a GelCount ^™^ Colony Counter (Oxford Optronix Ltd). The surviving fractions (SF) were calculated and normalized to the seeding efficiency. The survival curves were fitted in Prism 8 (GraphPad) using linear quadratic model: ln (SF) = - αD – βD^2^.

### Data Availability

The mass spectrometry proteomics data have been deposited to the ProteomeXchange Consortium via the PRIDE ^72^ partner repository with the data set identifier PXD018299.

