## Supporting Information for "Deep analysis of the USP18-dependent ISGylome and proteome unveils important roles for USP18 in tumour cell antigenicity and radiosensitivity"

**Running Title:** USP18 deletion enhances tumour cell immunogenicity and radiosensitivity

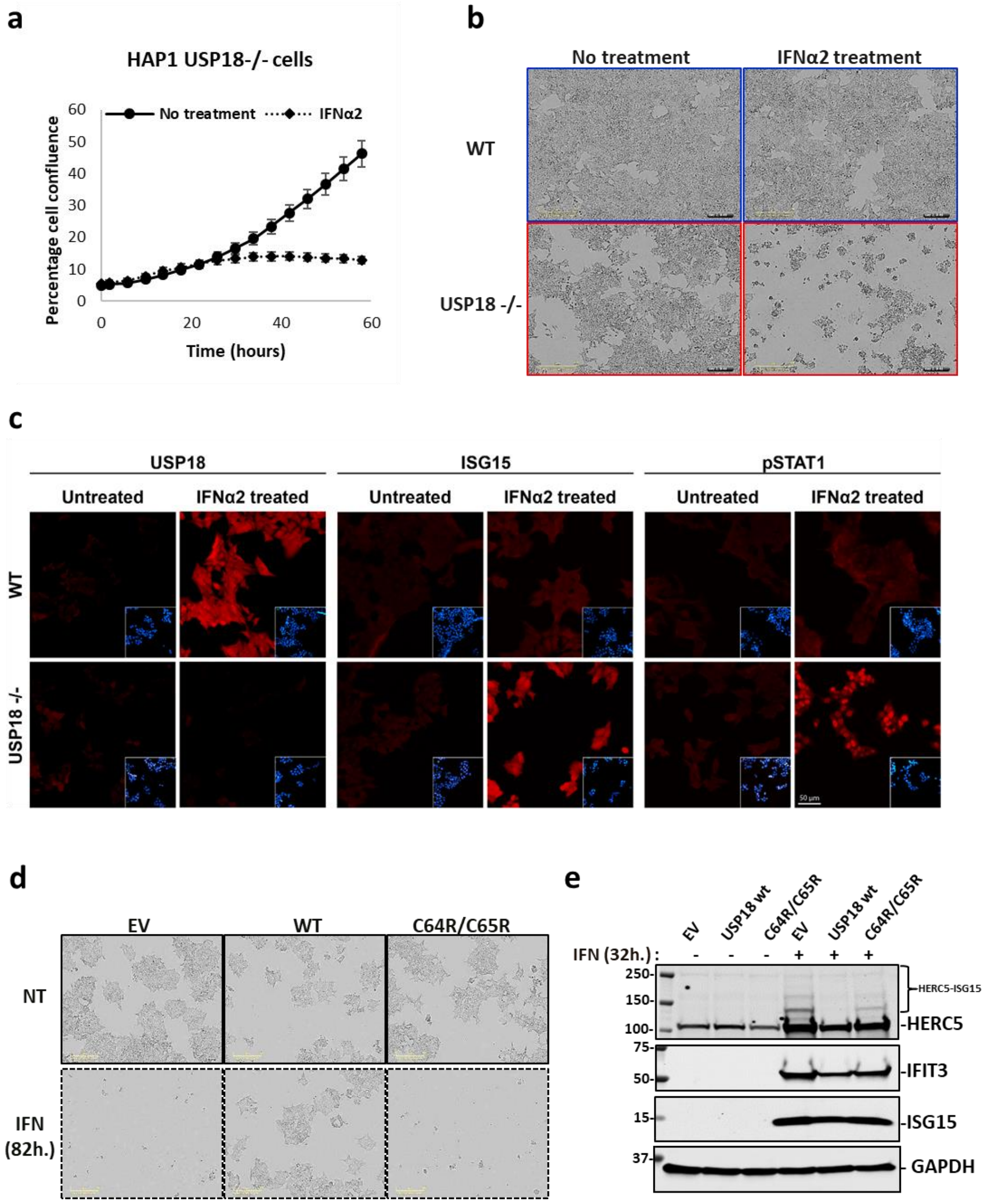


**Figure S1: USP18 alters protein ISGylation and cell viability in presence of IFN. a)** USP18-deficient HAP1 cell confluence (IncuCyte data) in the presence/absence of IFN treatment for 60 hours. **b)** Cell confluence and morphology of WT and USP18 KO cells after 58 hours of incubation with the indicated treatments (phase contrast images). **c)** Confocal microscopy immunofluorescence images displaying a strong accumulation of USP18 in WT cells, ISG15 and phosphorylated STAT1 (pSTAT1) in the USP18-deficient cells after 24 hours treatment with IFN. **d)** Cell confluence and morphology analysis showing that stable re-expression of USP18 WT, but not of a catalytically inactive mutant USP18 C64R/C65R, in USP18-deficient cells rescue them from the cytotoxic effects of IFN (82 hours treatment; EV: empty vector; NT: no treatment). **e)** Immunoblot of cell extracts from USP18 KO cells expressing USP18 WT and USP18 C64R/C65R. Ectopic expression of WT, and in a lesser extent catalytically inactive USP18, prevents IFN signalling as the expression of the ISGs HERC5, IFIT3 and ISG15 is reduced in these conditions. GAPDH was used as a loading control in the immunoblots (same experiment as in Figure 1g). Supporting data for Fig. 1.

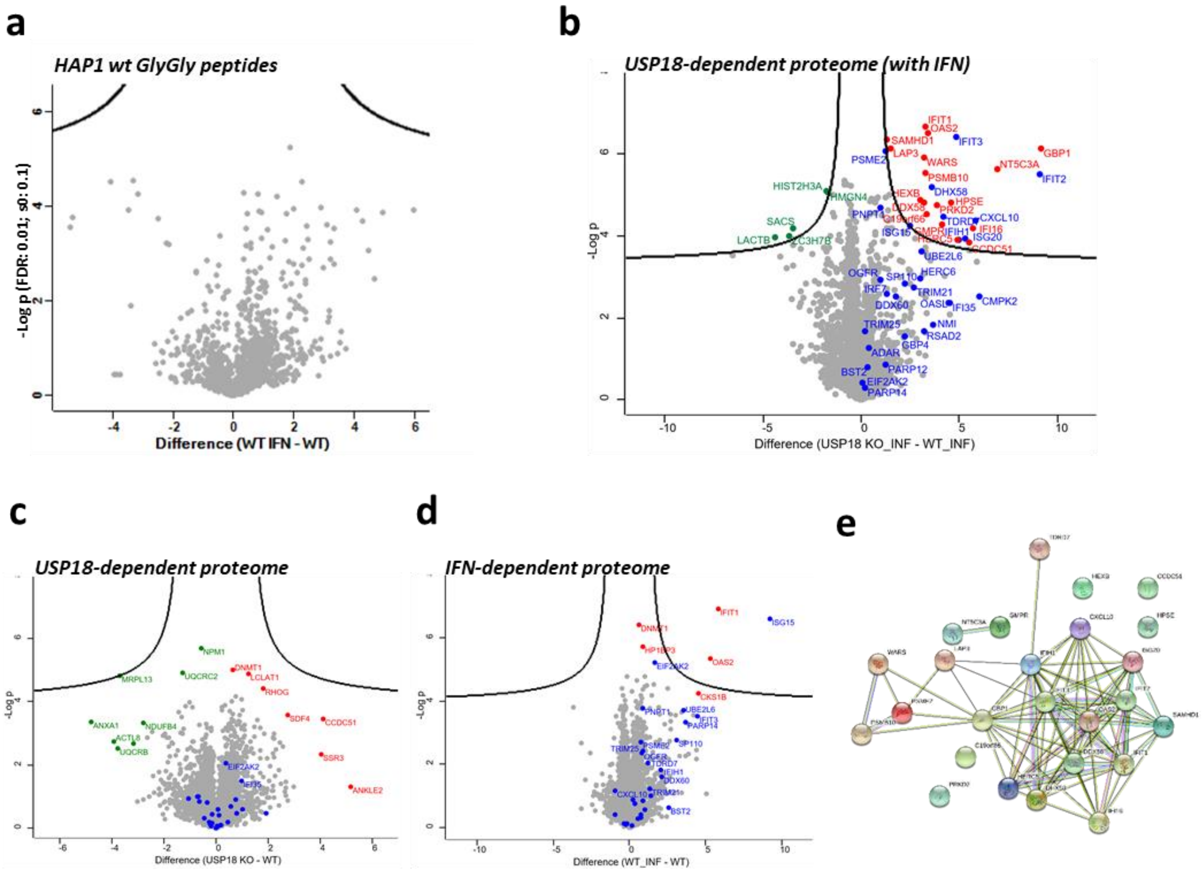


f

| source | term_name | term_id | adjusted_p_value | negative_log10_of_adjusted_p_value |
|---|---|---|---|---|
| GO:MF | catalytic activity, acting on RNA | GO:0140098 | 0.000172615 | 3.76292072 |
| GO:MF | RNA binding | GO:0003723 | 0.000465408 | 3.332166014 |
| GO:MF | double-stranded RNA binding | GO:0003725 | 0.000845981 | 3.072639239 |
| GO:MF | RNA helicase activity | GO:0003724 | 0.041074014 | 1.386432857 |
| GO:BP | defense response to virus | GO:0051607 | 1.92E-19 | 18.71633856 |
| GO:BP | response to virus | GO:0009615 | 1.83E-17 | 16.73868663 |
| GO:BP | defense response to other organism | GO:0098542 | 4.22E-12 | 11.37422576 |
| GO:BP | immune effector process | GO:0002252 | 1.19E-11 | 10.9237968 |
| GO:BP | innate immune response | GO:0045087 | 2.38E-10 | 9.624005607 |
| GO:CC | cytosol | GO:0005829 | 0.004816914 | 2.317231071 |
| KEGG | RIG-I-like receptor signaling pathway | KEGG:04622 | 0.000996999 | 3.001305297 |
| KEGG | Hepatitis C | KEGG:05160 | 0.02223447 | 1.652973213 |
| KEGG | Influenza A | KEGG:05164 | 0.029527012 | 1.529780498 |
| REAC | Interferon Signaling | REAC:R-HSA-913531 | 7.14E-09 | 8.146400099 |
| REAC | Immune System | REAC:R-HSA-168256 | 2.64E-07 | 6.579068538 |
| REAC | Interferon alpha/beta signaling | REAC:R-HSA-909733 | 5.03E-07 | 6.298493101 |
| REAC | Cytokine Signaling in Immune system | REAC:R-HSA-1280215 | 9.88E-07 | 6.005358344 |
| REAC | DDX58/IFIH1-mediated induction of interferon-alpha/beta | REAC:R-HSA-168928 | 0.003273145 | 2.485034747 |
| REAC | Antiviral mechanism by IFN-stimulated genes | REAC:R-HSA-1169410 | 0.003443526 | 2.462996681 |
| REAC | Negative regulators of DDX58/IFIH1 signaling | REAC:R-HSA-936440 | 0.008989984 | 2.046241084 |
| REAC | Innate Immune System | REAC:R-HSA-168249 | 0.020295503 | 1.692600172 |
| REAC | OAS antiviral response | REAC:R-HSA-8983711 | 0.032688155 | 1.485609594 |

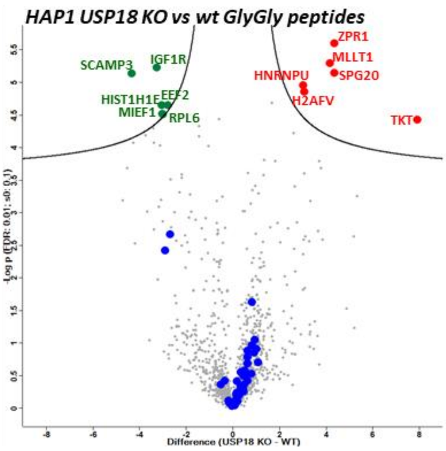


**Figure S2: Analysis of the USP18-dependent proteome. a)** Comparative volcano plot of the GlyGly modified peptides in HAP1 WT cells treated with IFN for 48 h. and untreated WT. No significant changes were observed using the statistical cut-off values used for all the proteomic analyses performed in this study (FDR: 0.01; s0: 0.1). Volcano plots showing the differential expression of proteins in the following comparisons: USP18 KO plus IFN vs WT plus IFN **b)**, USO18 KO vs WT **c)**, WT plus IFN vs WT **d)** (statistical parameters: FDR = 0.01; So = 0.1). STRING functional protein association networks analysis **e)** and g:PROFILER pathway enrichment analysis **f)** of the proteins signifantly up-regulated in A). **g)** Comparative volcano plot of the GlyGly modified peptides in HAP1 USP18 KO cells and WT cells (untreated).

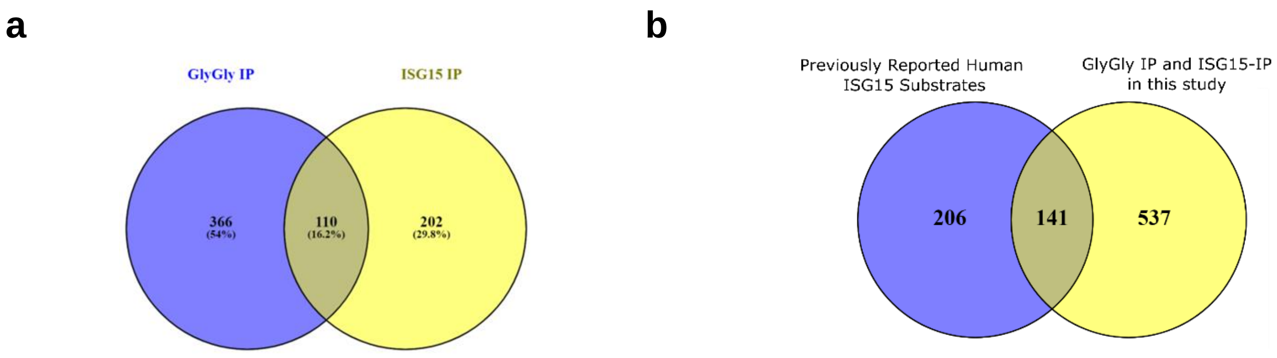


**Figure S3: USP18-deletion exacerbates the cellular ISGylated protein network. a)** Venn diagram showing the overlap between the GlyGly peptide IP and the ISG15 IP, performed in the same experimental conditions (HAP1 USP18 KO with IFN vs. HAP1 WT with IFN). **b)** Venn diagram comparing the depth of this ISGylome study and previously reported ISGylomes. Supporting data for Fig. 3.

| Up-regulated protein in melanoma patients responders to anti-PD1 | Proteome (USP18 KO_IFN vs WT_IFN) | GlyGly (USP18 KO_IFN vs WT_IFN) | ISG15 IP (USP18 KO_IFN vs WT_IFN) |
|---|---|---|---|
| PSME1 | UP (ns) | | DOWN |
| B2M | UP (ns) | | unchanged |
| TAPBP | UP (ns) | | |
| TAP2 | UP (ns) | | UP (ns) |
| CBR3 | | | UP (s) |
| TAP1 | UP (ns) | UP (s) | UP (s) |
| HLA-A | | | |
| HLA-C | | | UP (ns) |
| CD74 | | | |
| STAT1 | UP (ns) | UP (s) | UP (s) |
| IFIT1 | UP (s) | UP (s) | UP (s) |
| ICAM1 | | | |
| OGDH | unchanged | | unchanged |
| HLA-B | | | |
| HLA-DRA | | | |
| GBP1 | UP (s) | UP (s) | UP (s) |
| ERAP1 | UP (ns) | | UP (ns) |
| HLA-DPA1 | UP (ns) | | |
| IFI30 | UP (s) | | |
| HLA-DMB | UP (ns) | | |
| PSMB10 | UP (s) | UP (s) | |
| PSMB9 | UP (ns) | UP (s) | UP (s) |
| GBP2 | | UP (s) | |
| HMGCL | | | |
| HLA-DRB1 | | | |
| MAGEC2 | MAGED1/D2 (unchanged) | MAGED1 is UP (s) | MAGED2 is unchanged) and MAGED1 is UP (ns) |
| MX1 | | | MX2 is UP (s) |
| PSMB8 | DOWN | | |
| PARP14 | unchanged | UP (ns) | UP (s) |
| OPTN | UP (ns) | | |
| PARP4 | | | UP (s) |

**Supplementary Table 1:** Comparative between the up-regulated proteins in melanoma patients responding to anti-PD1 therapy [50] and our different USP18-dependent (USP18 KO cell plus IFN vs WT plus IFN) proteomic analysis datasets (Full proteome, GlyGly ISGylome and ISG15 interactome). (UP = up-regulated, DOWN = down-regulated, (s) = significant (t-test with permutation FDR = 0.01 for multiple-test correction and s0 = 0.1 as cut-off parameters) and (ns) = no significant.
